# Flap endonuclease Substrate Interactions studied using Dynamic Single-Molecule Atomic Force Microscopy

**DOI:** 10.1101/2024.01.10.574983

**Authors:** Vinny Verma, Emma L. Brudenell, Sophie Cussons, Janine Phipps, Su Chii Kong, Jon R. Sayers, Jamie K. Hobbs

**Affiliations:** Department of Physics and Astronomy, University of Sheffield (UK); Sheffield Institute for Nucleic Acids and Florey Institute, Section of Infection & Immunity, Division of Clinical Medicine School of Medicine and Population Health, The University of Sheffield, (UK); The Florey Institute for Host-Pathogen Interactions, University of Sheffield, UK; Sheffield Institute for Nucleic Acids, University of Sheffield, UK

**Keywords:** Atomic force microscopy, dynamic imaging, biophysics, flap endonuclease, mechanistic enzymology

## Abstract

Flap endonucleases (FENs) recognise and cleave DNA substrates containing a 5’-single-strand (ss) of nucleic acid branching off a double-stranded (ds) DNA to yield a nicked duplex during DNA replication. Dynamic Atomic Force Microscopy of an inactive FEN mutant complexed with branched DNA revealed mobilisation of immobilised DNA, indicating that protein interaction affected substrate conformation and disrupted the forces that anchored it to the poly-L-ornithine -treated mica surface. Enzymatically-active FEN was seen intermittently binding DNA, altering its conformation and cleaving the ssDNA branch. We developed a method using motion tracking for quantifying the movement of DNA sections, by visually segmenting DNA and tracking each segment to recognise the DNA sections most affected by the protein. It was found that whilst bound, FEN caused localised DNA bending, and changes in DNA shape were witnessed in the short time span of the protein’s appearance close to the nucleic acid, followed by protein adsorption on the mica surface. The results provide the first dynamic observations of FEN-DNA interaction. FEN initially binds to the dsDNA, slides to find the ds/ssDNA junction, and the 5’ ssDNA likely threads through a hole in the enzyme which leads to enzymatic hydrolysis of the branched substrate.

## Introduction

Flap endonuclease activity is required for processing of transiently branched nucleic-acid structures such as Okazaki fragments generated on the lagging strand during DNA replication^1^. In bacteria, FEN activity is provided by DNA polymerase I^2^, an enzyme which is comprised of two domains: the larger claw-shaped Klenow fragment and a smaller, N-terminal domain. This FEN-containing domain is responsible for nucleolytic cleavage of the 5’ ss-RNA branches on dsDNA during lagging-strand synthesis^3–7^. Some bacteria also carry an additional gene encoding a homologous protein consisting of a FEN domain, but no polymerase activity ^6,8,9^. A functional FEN domain is essential for cell viability in bacteria^10^. In eukaryotes and some bacteriophages, FEN activity is encoded by a discrete protein with no polymerase activity ^11,12^; Mice carrying only one functional FEN-encoding allele (*fen^+^*/*fen^-^*) develop tumours more rapidly than their *fen^+^*/*fen^+^* littermates, while homozygous knockout embryos die *in utero* suggesting that a FEN is essential in mice^13^.

Since Lyamichev and co-workers^14^ first suggested a threading mechanism could account for enzymatic cleavage at the junction between double and single-stranded DNA by bacterial DNA polymerase I, much research has been undertaken on recognition, binding and activity of FEN enzymes from eukaryotes^15–38^, prokaryotes ^6,8–10,39,40^ and bacteriophage enzymes ^12,36,41–50^,. These studies, coupled with crystal structures of diverse FEN-family enzymes ^35,51–54^, have identified critical residues forming the FEN active site. FEN substrate hydrolysis involves two divalent metal ions located in a cleft, capped with a loop or helical arch through which it was proposed that DNA might pass ^43–45,55,56^. Studies have led to the proposal of a mechanism of substrate recognition known as the ‘Threading model’ ^14^, (as shown in Figure 1) that explains the importance of the ssDNA branch passing through a hole in the protein to reach the active site. Here, divalent cations act as Lewis bases to catalyse phosphodiester bond cleavage yielding a ligatable nicked duplex and single-stranded nucleic acid^57^. A revised model proposes the binding of FEN to the branch point of DNA followed by bending of the ssDNA, threading and cleavage rather than the unlikely event of an enzyme first threading onto a free 5’ end ^21,40^.

**Figure 1:**
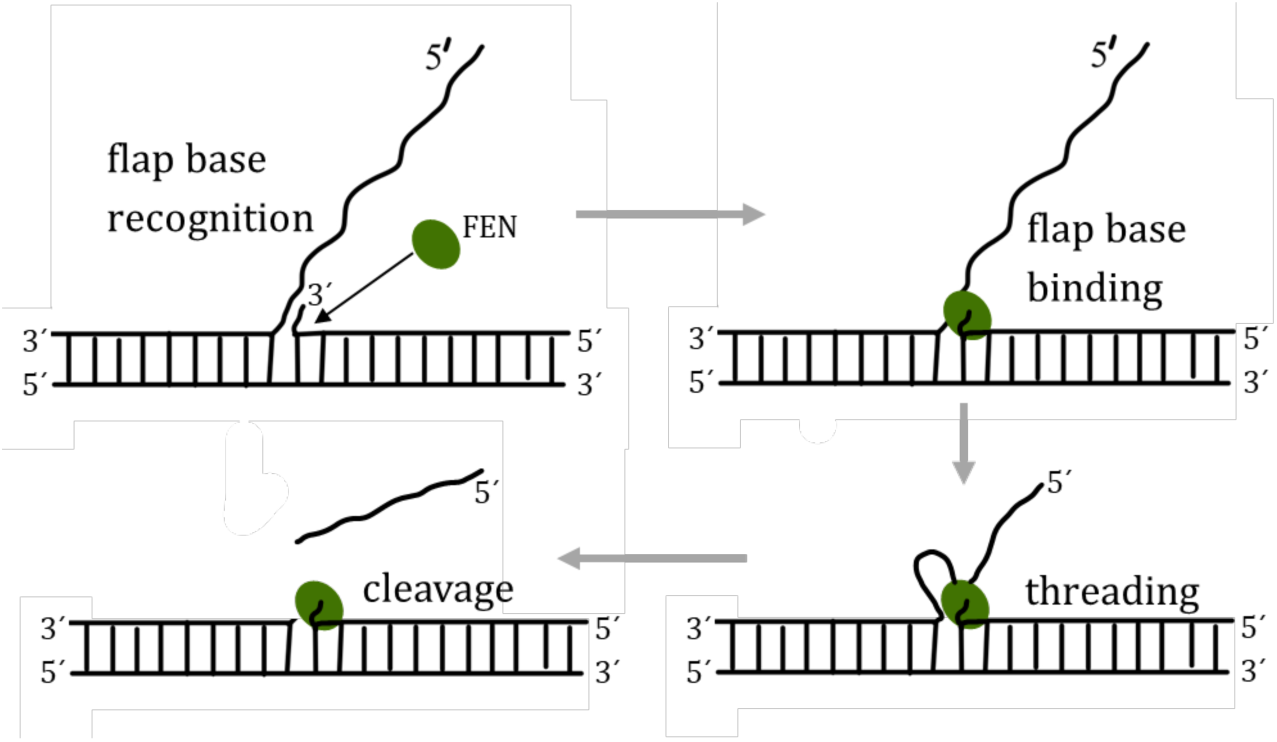
The ‘Threading model’ for flap DNA recognition and binding by FEN. ^21^: FEN binds the branch point, followed by bending and threading of the 5’ end and cleaving of the ssDNA. The diagram is not drawn to scale.

However, how the protein recognises the branched nucleic acid substrate in the cell in the first place is not well understood even though crystallography of bacteriophage and eukaryotic FEN homologues reveal that the 5’ single-stranded flap strand does indeed appear to thread through the enzyme^53,58^. A dynamic imaging technique that allows the visualization of the mechanism of substrate recognition and binding could enable an understanding of the process, complementing the information obtained from X-ray crystallography studies. Hence, in the current study, Atomic Force Microscopy (AFM) has been applied to visualize the FEN interaction with flap DNA to not only visualise the mechanism of binding and action of FEN, but also to formulate a standard method of nucleic acids-protein imaging and image data analysis to quantify the change in the conformation of nucleic acids due to their interaction with proteins.

Atomic Force Microscopy, invented by Binnig *et al* ^59^ as a type of scanning probe microscope, is an imaging technique that allows topographical mapping of conducting and insulating surfaces with nanometre resolution ^60,61^. For imaging with AFM, it is essential that the sample is immobilized on a flat surface to prevent any movement of the sample by the cantilever tip ^61^. This has been perfected over the years by different sample preparation methods in different laboratories ^62–71^. Since a DNA molecule has a negatively charged backbone, a positively charged surface can be used to bind the molecule with the aid of charge-charge interactions. Mica surfaces treated with divalent cations or synthetic amino acids such as poly-L-ornithine (PLO) and poly-L-lysine (PLL) have been used to immobilize DNA molecules for AFM imaging. While studying the binding and interaction of protein with DNA, it is important that the latter is immobilized on the surface strongly enough for the cantilever to detect it without any sample movement that might overly compromise the resolution of the images. At the same time, it is important that the DNA remains mobile enough to allow protein interaction.

Here, we report our observations of dynamic AFM imaging of flap DNA-FEN interactions carried out on a modified mica surface in buffer conditions with minimum sample damage or interference. A low concentration of the enzyme and divalent cations in the imaging buffer was used to control the rate of the processes observed during AFM visualization ^72^. In order to interpret these results, we developed a novel AFM image analysis pipeline to quantify DNA conformational changes due to protein interaction by molecule tracking and length measurements. Our results further our understanding of the structure specificity of the DNA - FEN interaction.

## Materials and Methods

Bacterial strains *E. coli* BL21^73^ and XL1-Blue^74^, were used for expression and amplification of pTTQ18 and derivatives, respectively. *E. coli* strain K12 ι1H1^75^ was used for pJONEX4 plasmid derivatives. Restriction endonucleases were supplied by New England BioLabs or Promega Corp. General laboratory chemicals were supplied by Sigma Aldrich, Merck or Thermofisher unless otherwise stated.

### Reagents for Sample Immobilization for AFM

Polyornithine, mica discs, CaCl_2_, KCl, MgCl_2_, NiCl_2_, HEPES (pH 7.8), HPLC water.

### Construction of expression plasmids used in this study

A vector containing a T5 promoter/lac operator sequence was constructed based on that present in plasmid pPANK ^76^ (nucleotides 1-87, GenBanK accession number AY327140). Briefly, oligonucleotides (sequences presented in Table S1) T5Pr02, T5Pr03, T5Pr05, T5Pr06, and T5Pr07 were individually phosphorylated, annealed together with oligonucleotides T5Pr01 and T5Pr04 then shotgun-ligated between the SapI and XbaI sites of plasmid pTTQ18^77^ using standard methods^78^. This cassette includes a T5 promoter with two *lac* repressor binding sites upstream of a translation initiation region incorporating an NdeI restriction site. The plasmid was designated pT5P (Figure S1 in Supplementary Materials).

A synthetic gene encoding a codon-optimised Taq polymerase mutant (Asp141Lys) was assembled from two synthetic fragments as follows: the N terminal FEN domain of Taq polymerase (with an Asp141Lys mutation (Bio Basic Inc, Canada)) flanked by an EcoRI site and ribosomal binding site (GAATTCTAAAAAGGAGGAAAACAT) with a 3’ BamHI site was used to replace the equivalent EcoRI-BamHI fragment in a pT5P carrying full-length Taq polymerase (supplied by Genscript). See Supplementary materials (Figure S2). Sequences of these gene fragments are provided in Supplementary Materials.

#### T*_7_* Exonuclease (a fully enzymatically active FEN homologue)

Bacteriophage T7 gene 6, encoding the exonuclease, was amplified from genomic DNA kindly supplied by Ian Molineux, University of Texas. Briefly, standard PCR amplification was carried out using primers T7exoF and T7exoR (Supplementary Table 1, which amplify residues 17467-18407 of Genbank entry LR745710) by 15 cycles of denaturation (95°C), 55°C annealing and 2 min extension times with Taq polymerase. The forward primer spanned an endogenous SnaBI site just upstream of the T7 gene 6 Shine Dalgarno region. The reverse primer introduced a TAA stop codon to the end of the T7 gene 6 coding region followed immediately by a PstI restriction endonuclease recognition site. The resulting PCR product was purified by agarose gel electrophoresis and recovered using a Promega Wizard Gel clean-up kit according to the manufacturer’s instructions. This fragment was cut with SnaBI and PstI ligated into Eco53KI/PstI double-digested plasmid pJONEX4 and the ligation products transfected into K12 ΔH1 *E. coli* grown at 28°C on 2yT agar plates supplemented with 100 ug/mL ampicillin. Successful cloning was confirmed by restriction analysis and DNA sequencing (University of Sheffield Core Genomics facility) and the resulting plasmid designated as pJONEXT7exo.

### Protein Production

The bacteriophage T7 exonuclease (T7FEN) was overexpressed in *E. coli* ΔH1 from plasmid pJONEX4 under the control of the heat-inducible λP_L_^79^. Transformed cells were grown at 28°C until mid-log phase and protein overexpression was induced at 42°C for 2 hours. Cells were harvested immediately after induction by centrifugation at 12,000 x *g* for 30 minutes and pellets stored at -80°C.

All stages of protein purification were carried out at 4°C. Cells were resuspended in 5 mL/g lysis buffer (25 mM Tris-HCl pH 8, 100 mM NaCl, 5% (v/v) glycerol, 2 mM EDTA, 1 mM DTT, 10 µg/mL 4-(2-aminoethyl)benzene sulfonyl fluoride hydrochloride(AEBSF)) and lysed by the addition lysozyme (1 mg/mL) and sodium deoxycholate (500 µg/mL) until lysed. Viscosity was reduced by subjecting the lysate to ∼ 30 s ultrasound bursts in an MSE Soniprep 150 Plus sonicator until the lysate became free flowing. Nucleic acid was removed from soluble lysate via precipitation with 0.35% (v/v) polyethyleneimine (PEI.HCl, pH8). Residual PEI was removed by precipitation of protein by addition of solid ammonium sulphate to a final concentration of 3.5 M. The precipitate was resuspended and dialysed overnight twice into 20-fold excess KP7/50 buffer (25 mM potassium phosphate pH 7, 5 % (v/v) glycerol, 2 mM EDTA, 1 mM DTT). Soluble lysate was loaded onto a prepacked 20 mL HiTrap Heparin HP column and protein was eluted over a gradient of buffer containing 50 mM -1 M NaCl. Protein was purified further using Q Sepharose HP in Tris/50 (25 mM Tris-HCl pH 8, 5% (v/v) glycerol, 2mM EDTA, 1 mM DTT) and SP Sepharose by elution with a 50 mM – 1 M NaCl gradient in KP7/50 buffer. Protein was concentrated and purified further by size exclusion chromatography using a S200 16/60 column (GeHealthcare). Peak fractions were concentrated and stored at -20°C in 50% (v/v) glycerol. Protein purity was assessed by SDS-PAGE and intact protein mass was determined using Matrix-assisted Laser Desorption/Ionisation time-of-flight (MALDI-TOF) (figure 4) by the University of Sheffield Mass Spectrometry service (Department of Chemistry, UoS).

#### *Thermus aquaticus* DNA Polymerase I (Asp141Lys)

Recombinant BL21 cells carrying plasmid TaqPolAsp141Lys were fermented, lysed using lysozyme in the presence of ethylenediaminetetraacetic acid (EDTA), sodium deoxycholic acid and phenylmethylsulphonyl fluoride (PMSF), incubated 37℃/1 hr and sonicated to reduce viscosity and release soluble proteins ^80,81^. The cell debris was removed by centrifugation at ∼40,000 x *g* for 20 min. The supernatant was removed and pasteurized (70℃ for 30 min) and the insoluble protein aggregates removed by centrifugation as above. The pellet was discarded and the supernatant treated with polyethyleneimine (PEI.HCl pH 8) to precipitate nucleic acids which were removed by centrifugation as above, followed by ammonium sulphate (3.5 M) precipitation ^81–85^ of protein to remove PEI. The protein pellet was suspended in 50 mM salt/EDTA/25 mM phosphate buffer (pH 8) with 5% glycerol. This solution was dialysed against the same buffer overnight, to remove the ammonium sulphate.

Anion exchange chromatography on a prepacked 20 mL HiTrap Heparin HP column across salt gradient 50 mM to 1 M was used to separate the TaqPolAsp141Lys from contaminating proteins. The combined fractions of TaqPolAsp141Lys were dialysed against 50 mM salt/EDTA/25 mM TRIS buffer overnight and purified on the Q Sepharose HP column across salt gradient 50 mM to 1M. The protein was further purified on the SP Sepharose column in 50 mM salt/EDTA/250 mM HEPES buffer across salt gradient 50 mM to 1M. To remove the impurities still present, ammonium sulphate precipitation at 1 M – 4 M concentrations of the salt was performed. Additional purification was carried out on SP Sepharose in pH 6 HEPES buffer and on a Q Sepharose HP column with a TRIS buffer (pH 9). Zymogram (see supplementary material for details of the assay) and UV nuclease assay were performed to ensure that the inactive FEN domain did not have any nuclease activity (Figure 3). Equal volume of glycerol and 0.1% sodium azide were added into the samples to flash freeze in liquid nitrogen. The samples were then stored at -80℃.

**Figure 2:**
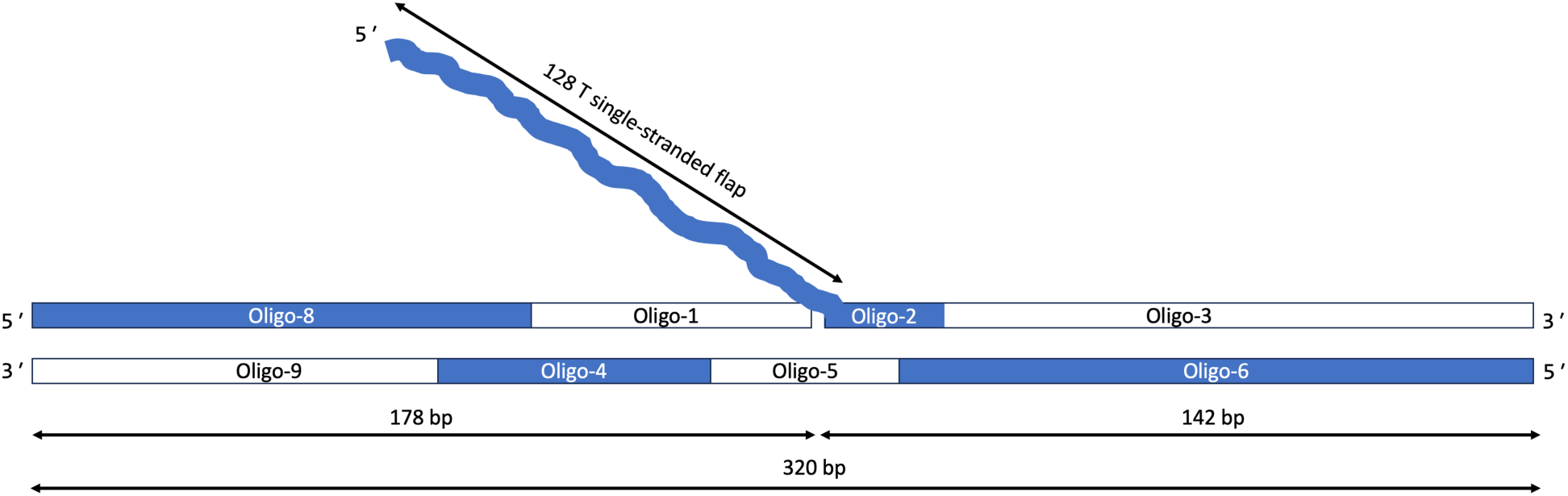
**Schematic of flap DNA formed by oligonucleotide assembly and observed using AFM. The details of the Oligo 1-9 in Supplementary Information Table S1.**

**Figure 3:**
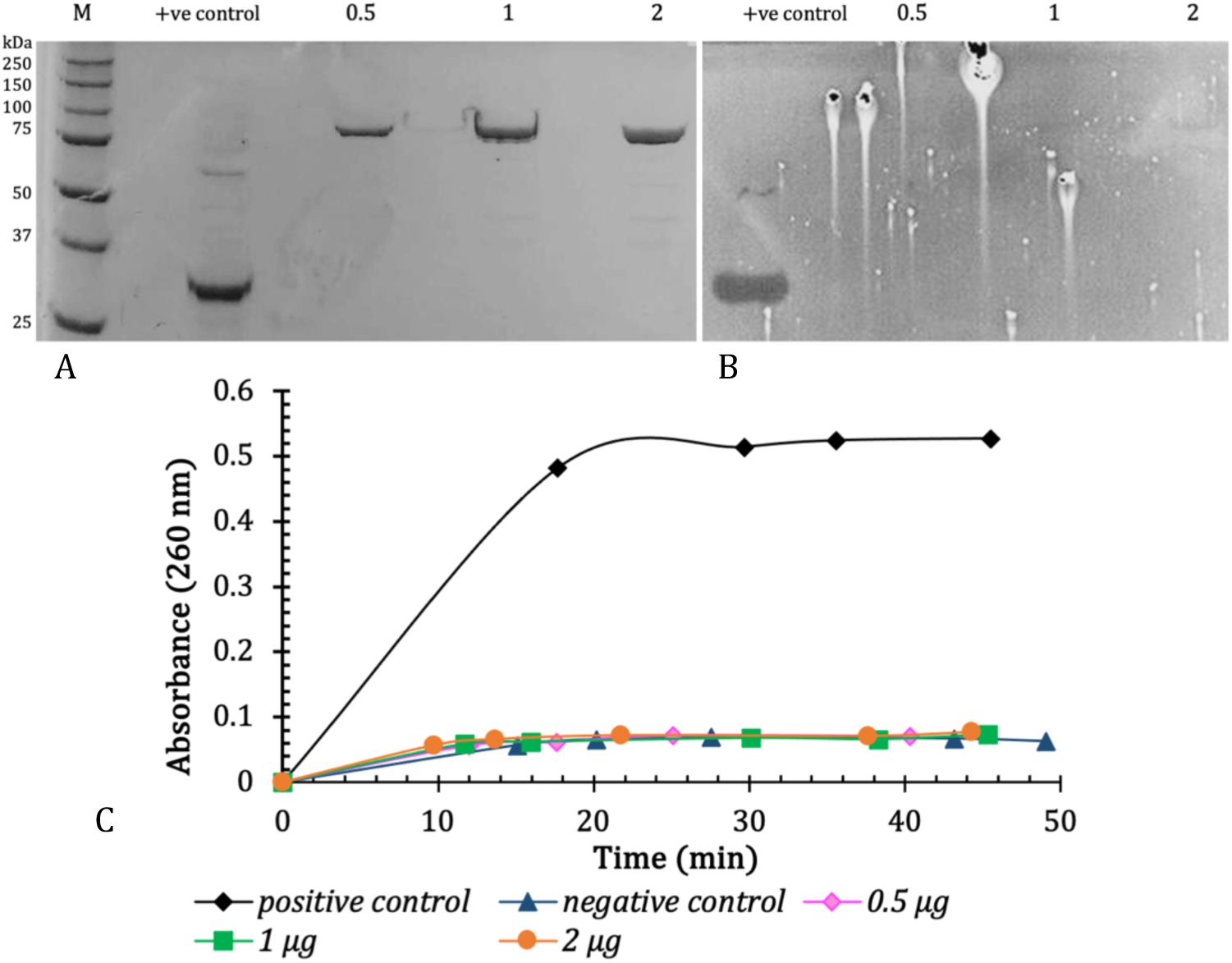
Analysis of purified TaqPolAsp141Lys. (A) SDS-PAGE, (B) zymogram and (C) UV-Nuclease assay. +ve control-T7 nuclease: 1.5 µg in (A) and (B), 0.5 µg in (C). Negative control in (C) is MQ-H_2_O. The zymogram gel was incubated in reaction buffer for 30 minutes. Lane numbers denote amount of protein loaded in μg. Analysis was performed on 10% acrylamide gels with PrecisionPlus marker (BioRad) (lane M) used.

**Figure 4:**
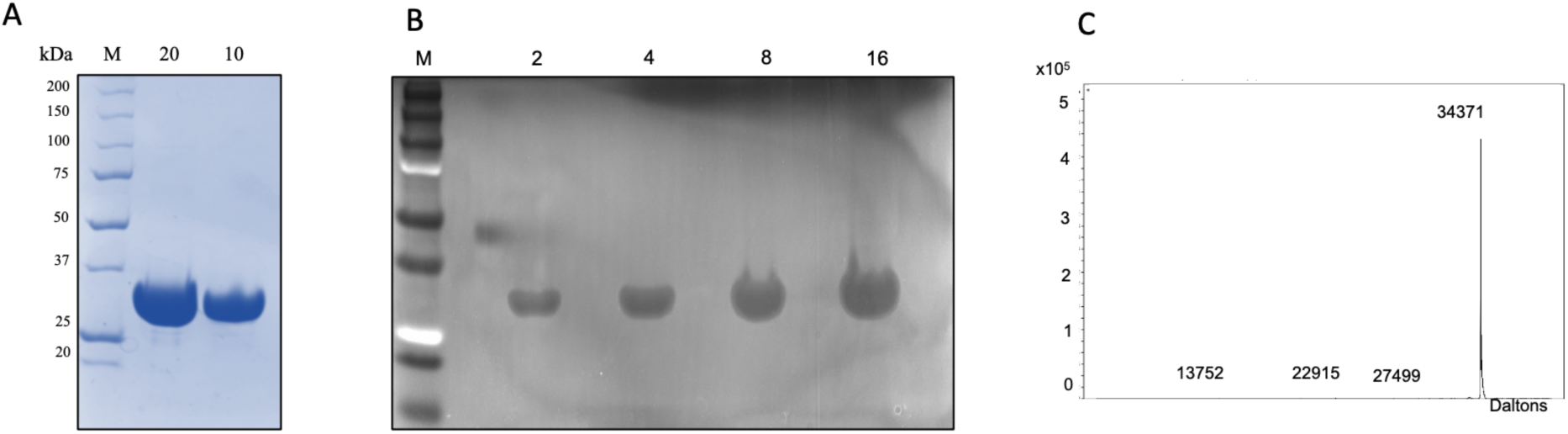
Analysis of purified T7 exonuclease. Purified T7 exonuclease was analysed by **(A)** SDS-PAGE, **(B)** zymogram and **(C)** MALDI-TOF mass spectrometry. The zymogram gel was incubated in reaction buffer for 15 minutes. Lane numbers denote amount of protein loaded in μg. Analysis was performed on 10% acrylamide gels with PrecisionPlus marker (BioRad) (lane M) used.

### Production of 320 bp flap DNA

Oligonucleotides of varying lengths were designed and ordered from Eurofins such that they could be assembled to form flap DNA of length 320 bp with a ssDNA flap of length 128 nt (see Figure 2 and Table S1 for details of the process and oligonucleotide sequences). Ligation and annealing of the oligos performed by preparation of 25 µL an aqueous mix of 2.5 µL T4 DNA ligase buffer (10X), 1 µL T4 DNA Ligase (NEB), 100 ng DNA (oligo 3), 2.5 µL PEG 4000 (5%). The reaction mix was incubated at 37°C for 30 minutes followed by heat inactivation of the enzyme at 65°C for 20 minutes. The mixture was then cooled on ice. The concentration of the DNA was estimated by UV spectrophotometry.

### AFM experimental conditions

Flap structures were assembled by annealing the constituent synthetic oligonucleotides and ligation *in vitro* as described in Materials & Methods.

Bruker Fastscan Bio was used with Bruker TESPA-V2 (nominal tip radius 7 nm, nominal spring constant 37 Nm^-1^) and Nunano SCOUT 350 (nominal tip radius 5 nm, nominal spring constant 42 Nm^-1^) probes for imaging in air and Bruker Fastscan D (nominal tip radius 5 nm, nominal spring constant 0.25 Nm^-1^) probes for imaging in buffer. For dynamic imaging, DNA solution prepared in HEPES buffer (25 mM HEPES, pH ∼7.5) containing 2 mM Ca^2+^ and 5 mM K^+^ was immobilized on PLO-treated mica, washed with HPLC water and dried under N_2_. To image the interaction with inactive FEN, the sample prepared above was imaged in the same buffer as the DNA solution. In case of interaction with active FEN, the same buffer solution but with Mg^2+^ in place of Ca^2+^ and an additional 10 mM DTT was used for imaging. During imaging, protein solution (in HEPES buffer) was added (∼10-20 µL) onto the sample with caution to not disturb the imaging. Continuous images were taken to observe the conformational changes. The frame rate was usually limited to approximately 30 s per frame. 400×400 nm^2^ images were collected which were cropped to focus on the molecules of interest during image post processing. The frames were flattened (1^st^ order) using Bruker Nanoscope Analysis software and aligned to reduce the horizontal drift in the images followed by conversion to videos using Adobe Photoshop.

#### DNA Motion Tracking

A method of tracking the motion of DNA under the influence of protein was developed. Adobe Illustrator was used to mark segments of 5 pixel (or 10 pixels in the images where DNA was longer so reducing the number of segments and make the plots less crowded) on the DNA molecule in each of the frames. Each of the DNA segment were differently coloured to be tracked independently without confusion. These frames were then assembled into a video using Adobe Photoshop (by creating Time-lapse). The software ‘Tracker: Video Analysis and Modelling Tool’ was used to track each of the DNA segments and their respective displacements for each frame were plotted using Microsoft Excel. It should be emphasized that the displacements of each of the segments was measured as a magnitude of displacement vector from the previous position, and **<u>not</u>** from the initial starting position. The plots allowed us to track which section of the molecule moved and when, with respect to the position and time of the protein’s appearance in the vicinity of the DNA molecule (see Figure S3 for details of the process).

## Results and Discussion

The purified proteins TaqPolAsp141Lys and T7 exonuclease were analysed for activity by zymogram (see supplementary material for details of the assay) and UV nuclease assay (figures 3 and 4).

### Static high-resolution AFM images of flap DNA depict the large difference in the width of dsDNA vs ssDNA flap

As shown in Figure 5C, AFM images of flap DNA in HEPES buffer (pH 8) appear as a ssDNA branch attached on a dsDNA. The ssDNA is a narrower and lower strand on the AFM image (corresponding to the smaller width of the ssDNA) and the dsDNA is the taller strand, signifying its comparatively larger width.

**Figure 5:**
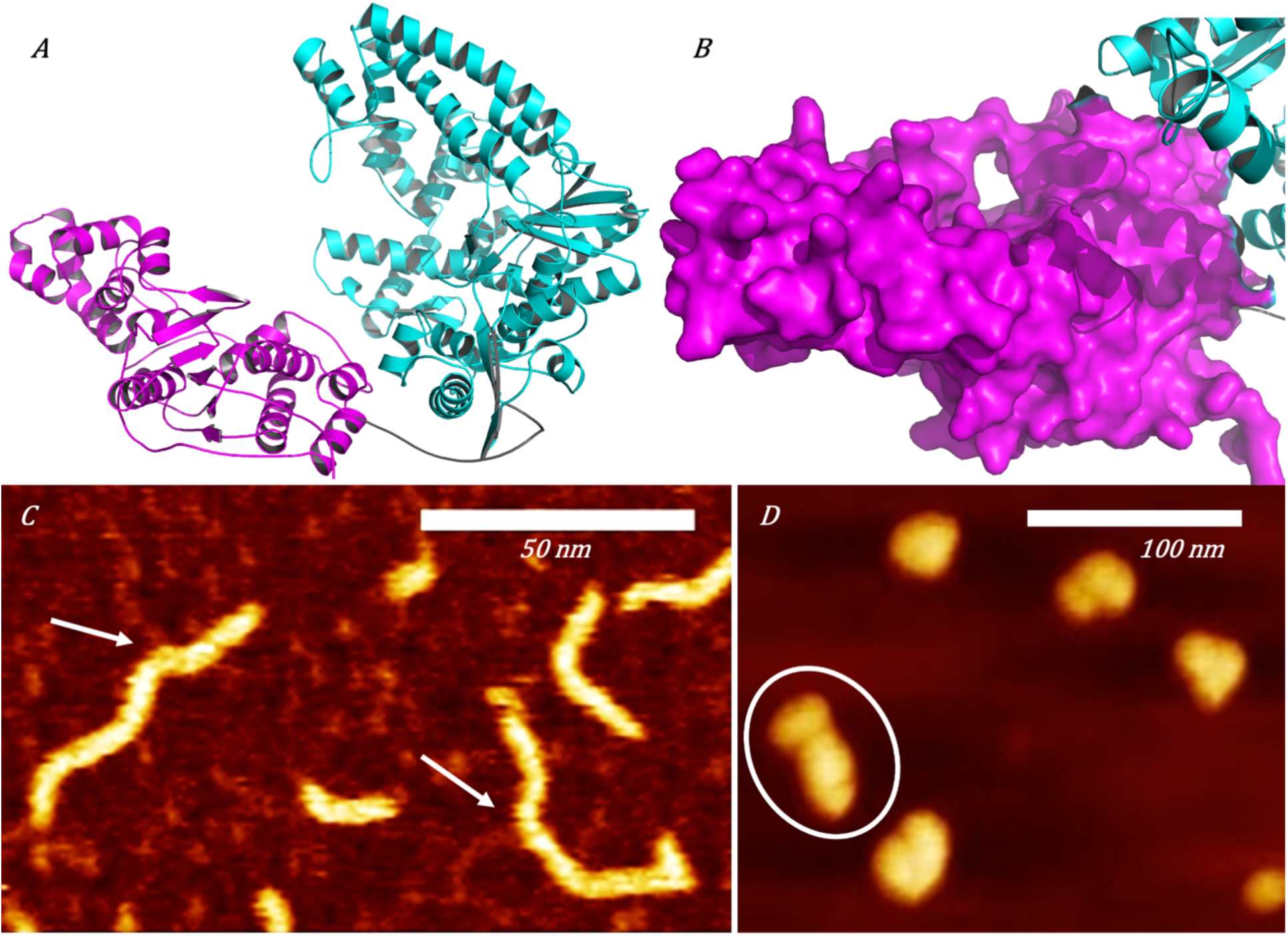
AFM images of flap DNA and TaqPolI (inactive FEN domain TaqPolAsp141Lys): **(A)** Cartoon representation of the structure of *Thermus aquaticus* DNA Polymerase I holoenzyme from AlphaFold (UNIPROT P19821). The Stoffel fragment shown in cyan (equivalent to the Klenow in the *E. coli* homolog) ^86^ with characteristic ‘hand, fingers and thumb’ structure is attached via a flexible linker (grey) to the FEN domain (magenta). **(B)** Enlarged view of the FEN domain shown with magenta surface. A hole thought to allow passage of single-stranded nucleic acid resides above the globular domain. **(C)** 320 bp flap DNA with 128 nt flap on PLO, imaged in buffer: height 4.5 nm. **(D)** TaqPolI immobilized on mica, imaged in air. Height 9 nm. Out of the 5 protein molecules shown here, one of the molecules shows the two domains resolved with the claw shape of the Klenow fragment distinguishable (encircled). The other molecules appear globular in air.

### Static AFM images of TaqPolI

As depicted from Figure 5B, TaqPolI in air appears in clumped conformations with some instances of elongated shapes seen (encircled in figure 5D), where the two domains of the TaqPolI were resolved. In these very rare cases, the claw shape of the Klenow fragment and the somewhat-oval shape of the FEN domain could be seen corresponding to the crystal structure of the molecule (Figure 5A and 5B).

### FEN modifies the electrostatic environment around DNA for binding

DNA Polymerase requires the presence of Mg^2+^ as a cofactor in the buffer at a suitable pH 8. Mg^2+^ allow efficient binding and activity of FEN domain of DNA PolI to DNA for the replication process to occur ^41,45,87–89^, while Ca^2+^ augment the binding of FEN alone to the DNA, without allowing any endonuclease activity ^47,89,90^. For this reason, inactive FEN was imaged in a buffer containing Ca^2+,^ to ensure protein binding, but DNA PolI containing active FEN domain was imaged in a buffer containing Mg^2+^ to observe FEN activity.

From the dynamic AFM images (figures 6A and 7A), we have observed that the presence of FEN (whether inactive or active FEN domain) appears to be capable of modifying the charge environment in the vicinity of DNA molecules causing dissociation from the PLO treated mica surface. DNA molecules are well immobilized on the PLO treated mica surface by electrostatic interactions during the consecutive scans before the addition of protein, as well as the negative control experiments of DNA imaged without the presence FEN or DNA PolI (*e.g.* frames 1-16 of Figure 6A and C, frames 1-9 of Figure 7A and C). However, FEN caused movement and change in the shape of DNA (most prominently seen between frames 18-19 and Figure 6A), either by binding onto DNA or on the PLO surface slightly displacing the DNA molecule during immobilization. The other DNA substrates distant from FEN molecules showed no change. Additionally, the motion of the section of DNA closest to the protein can be seen in the immediate time frames when the protein was first seen interacting with the DNA. The DNA appeared to be well immobilized until the protein first appeared close to the DNA (frame 18 in Figure 6A). From then on, the shape of the DNA changed indicating that the protein had bound to DNA causing the latter to come off the surface, move and immobilize again in a different conformation. In some cases, the DNA molecule appeared to bend at the point of contact with the protein (frames 9-12 in Figure 7A) and the protein also moved around the axis of the DNA molecule. A schematic of the DNA movement (Figure 6C and 7C) depicts the change in its conformation due to this interaction.

**Figure 6:**
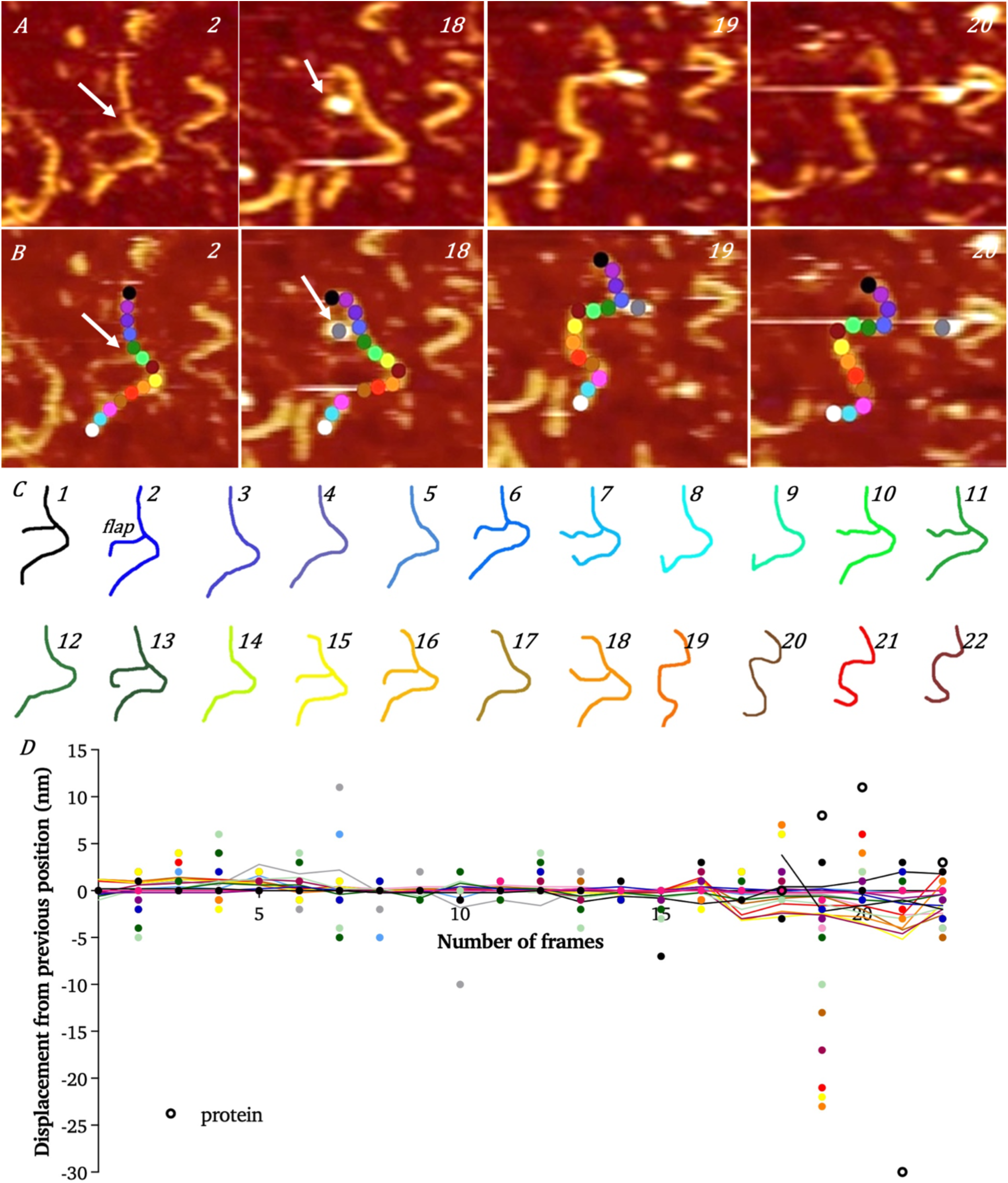
Flap DNA–inactive FEN domain TaqPolAsp141Lys imaged in buffer (25 mM HEPES, 2 mM Ca^2+^, 5 mM K^+^) immobilized on PLO treated mica, dynamic imaging of interaction with 320 bp flap DNA. **(A)** height 5 nm, scale bar 50 nm. The frames shown here are 2, 18-20 of a 22-frame scan. Refer to Movie A in Supplementary Material. Flap DNA seen in frame 2 shows a ssDNA flap (marked by arrow). The DNA shape changes because of protein binding. **(B)** Trace of movement of segments of DNA from figure (A). The DNA is labelled with markers of 5-pixel length. An aliquot of protein solution was added during frame 16 and immediately the DNA conformation changed in the consecutive frames 18-19. The protein is labelled as grey point appearing first in frame 18. **(C)** Sketch of DNA in the various frames. **(D)** 0^th^ order smooth (Savitzky and Golay smooth using least squares^91^) of displacement of segments of DNA in the presence of inactive FEN protein. The displacement lines of all the fragments/markers appear to be mostly synchronised until frame 18, with some markers showing slight deviations, namely the pink and blue markers around frames 6-11, that indicated the end of the DNA. These fluctuations were possibly due to the end of the DNA being more mobile.

**Figure 7:**
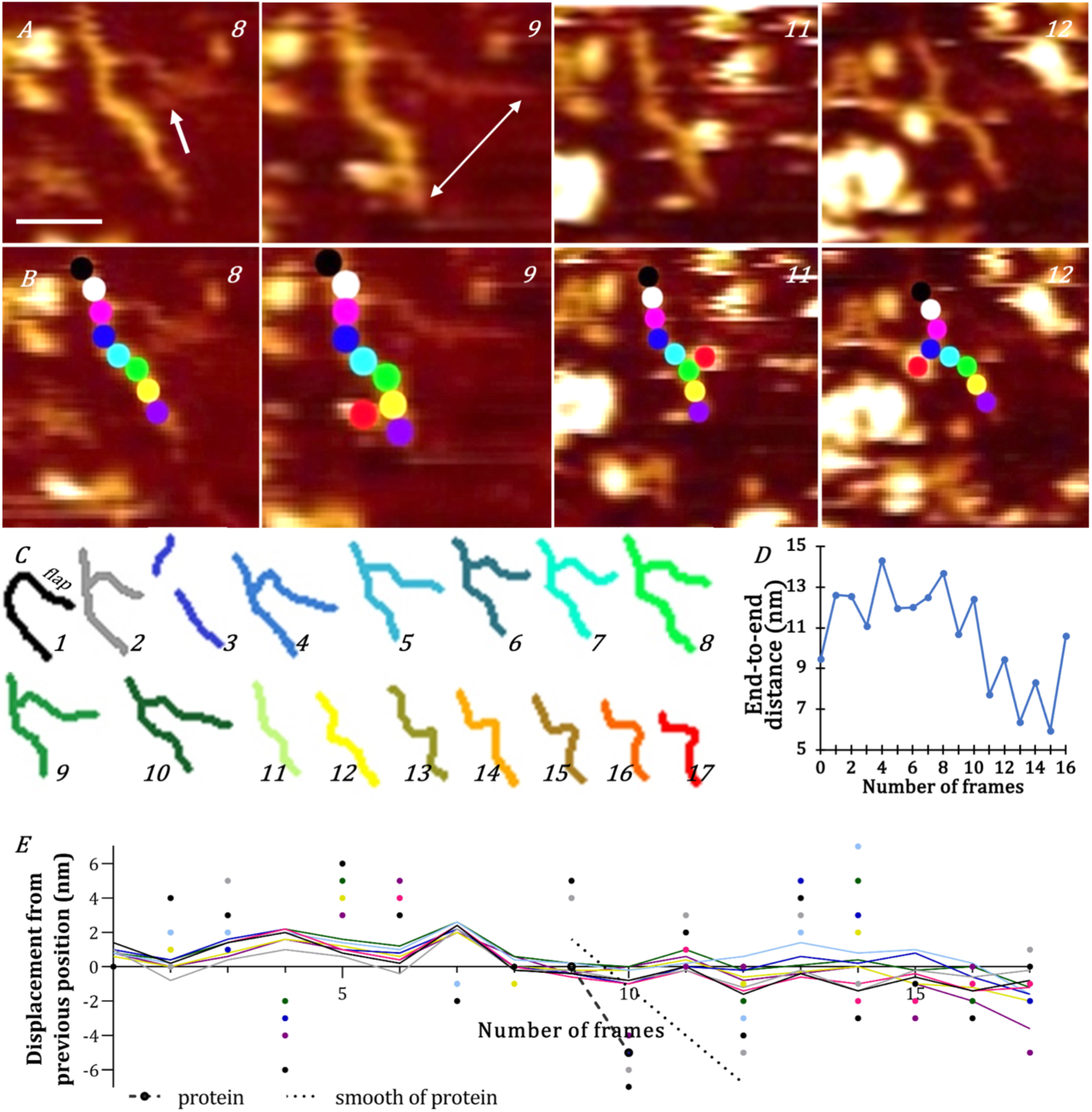
Flap DNA-inactive FEN domain TaqPolAsp141Lys) imaged in buffer (25 mM HEPES, 2 mM Ca^2+^, 5 mM K^+^) immobilized on PLO treated mica, dynamic imaging of interaction with flap DNA. **(A)** height 5 nm, scale bar 20 nm. The frames shown here are from a 17-frame scan. The protein was added in frame 3 and appeared about 3 minutes later near the DNA in frame 9. Refer Movie B in Supplementary Material. **(B)** Trace of movement of 10-pixel segments of DNA from figure (A). **(C)** Sketch of DNA in the various frames. **(D)** End-to-end distance of DNA in the presence of inactive FEN protein (the length measured is shown by arrow in frame 9 of figure A). The end-to end distance decreases as the protein appears close to the DNA frame 9 onwards. **(E)** Displacement of segments of DNA in the presence of inactive FEN protein.

We have quantified this motion by tracking DNA fragments (obtained by dividing the entire DNA strand into 10 nm segments, as shown in figures 6B and 7B) and plotting the 0^th^ order smooth (Savitzky and Golay smooth using least squares^91^) of their displacement-time plot or the corresponding motion (figures 6D and 7E). The displacement time plots appear to be synchronized and denote the well-immobilized DNA molecules. However, a significant fluctuation in the plots could be seen in frames 18-22 (Figure 6D) where the marker displacements were not only large but significantly unsynchronized, indicating the movement and conformational change of the DNA molecule. This indicated that the protein bound to the DNA causing some mobilization of the latter in the process. In some cases, the protein bound to DNA (in frame 9 of Figure 7A) caused a slight bend to be seen in the DNA, but did not produce any numerically significant deviation (Figure 7E). The plot however, indicated slight movement of some segments of DNA over frames 1-3, supported by the absence of synchronization of the plots during those frames. The presence of the protein from frame 9 onwards caused the DNA to move and disturbed the otherwise well-synchronized displacement plots over frames 11-17. The DNA moves between frames 13-14 and can be seen as a lack of synchronization of the plot lines of the DNA displacements. A measurement of the end-to-end distance of the DNA molecules (Figure 7D) also depicts the change in the DNA lengths in the presence of the protein and signifies the effect of the latter on the binding ability of DNA to the surface. These observations imply that protein binding to DNA causes a change in the electrostatic environment around the substrate. This affects DNA mobility and thus, its conformation, especially near the position of protein binding. Most likely the surface charges of the DNA Polymerase I or FEN interacted with DNA causing the latter to overcome the attractive forces of PLO binding it to the mica surface, loosen from the surface to facilitate protein binding, move (slightly or a lot, depending upon how easily the ssDNA can anchor the PLO treated mica surface) and immobilize again. In many instances, the protein either did not immobilize around the DNA or immobilized on the PLO surface close to the DNA giving the false impression that the protein had bound the DNA substrate. In those cases, no changes in the DNA conformation were observed.

### DNA polymerase I or FEN protein can bind anywhere on the DNA

From the previously reported crystallography studies, it is proposed that the protein is able to bind the DNA at the branch point of flap DNA, bend the 5’-single-stranded end towards it thus promoting threading through the protein arch (see figure 1). This positions the scissile phosphodiester bond in proximity to the active site divalent cation for catalysis ^21,43–45,55,56,58^. However, in the current AFM experiments, we have found that the protein (entire DNA Pol I or FEN alone) is able to bind the DNA anywhere (as marked by arrows in frame 18 of Figure 6A and frame 8 of Figure 7A), and not just at the point of attachment of the ssDNA branch to the dsDNA. The binding of the protein could be supported by the mobility of DNA in the vicinity and time frames of the protein’s presence next to it. Any random immobilization of the protein close to the DNA did not cause an observable change in the conformation of the immobilized DNA molecule, as depicted by the displacement frame plots figure 6(D) and 7(E). This indicates that FEN protein or FEN domain containing DNA PolI was capable of binding to the DNA anywhere, rather than specifically recognising the branch point at which FEN catalysis occurs. In the cases where the FEN *appeared* in proximity to the ssDNA end without actual binding, no movement of the ssDNA was apparent. No nuclease activity occurs when FEN was not bound close to the branch point but at the farther positions on the DNA as expected for a flap-structure specific endonuclease.

### Protein slides to the branch point

The dynamic images of inactive FEN-domain-containing TaqPolI interacting with flap DNA (frames 9-12 of Figure 7A and B) show that the protein is adept at sliding along the length of its substrate and bending it along its way. Though the DNA appears well immobilized, the impact of FEN binding is sufficient to overcome the electrostatic forces anchoring it to the surface. These observations suggest that FEN or DNA PolI containing FEN, after binding anywhere on the dsDNA, has the ability to ‘search’ the length of DNA for the bifurcation and consequently localise to the branch point where the scissile phosphodiester bond is located. Similar behaviour has been reported for other nucleic acid metabolising proteins ^92,93^. This fits with the ability of DNA Polymerases to bind DNA randomly and ‘scan’ for suitably primed replication sites but is contrary to earlier proposals of binding directly at the DNA branch point ^21,40^.

### FEN binding to the ssDNA observed using dynamic AFM imaging

Although AFM does not have the resolution to directly view threading in this system, we have imaged the proposed threading of ssDNA through FEN (Figure 8A), where FEN binding to the overhang DNA branch point caused the ssDNA end to bend towards it and thread through the protein. The end ‘a’ of the DNA can be seen breaking off the dsDNA end in frame 21-24, bends towards the protein (shown by blue arrow) in frame 37 and tries to bind to it in frame 38, and again in frames 43 and 48, after failing to latch onto it in frame 42. Once it is attached to the protein, the ssDNA end threads through the protein in frames 48 onwards, which supports the proposed concept of threading of ssDNA during the process of FEN catalysis. Interestingly, the ssDNA end did not bend and bind FEN immediately but flapped around before binding and became successful after multiple attempts. This change in the conformation can be visualized from the schematic Figure 8(C).

**Figure 8:**
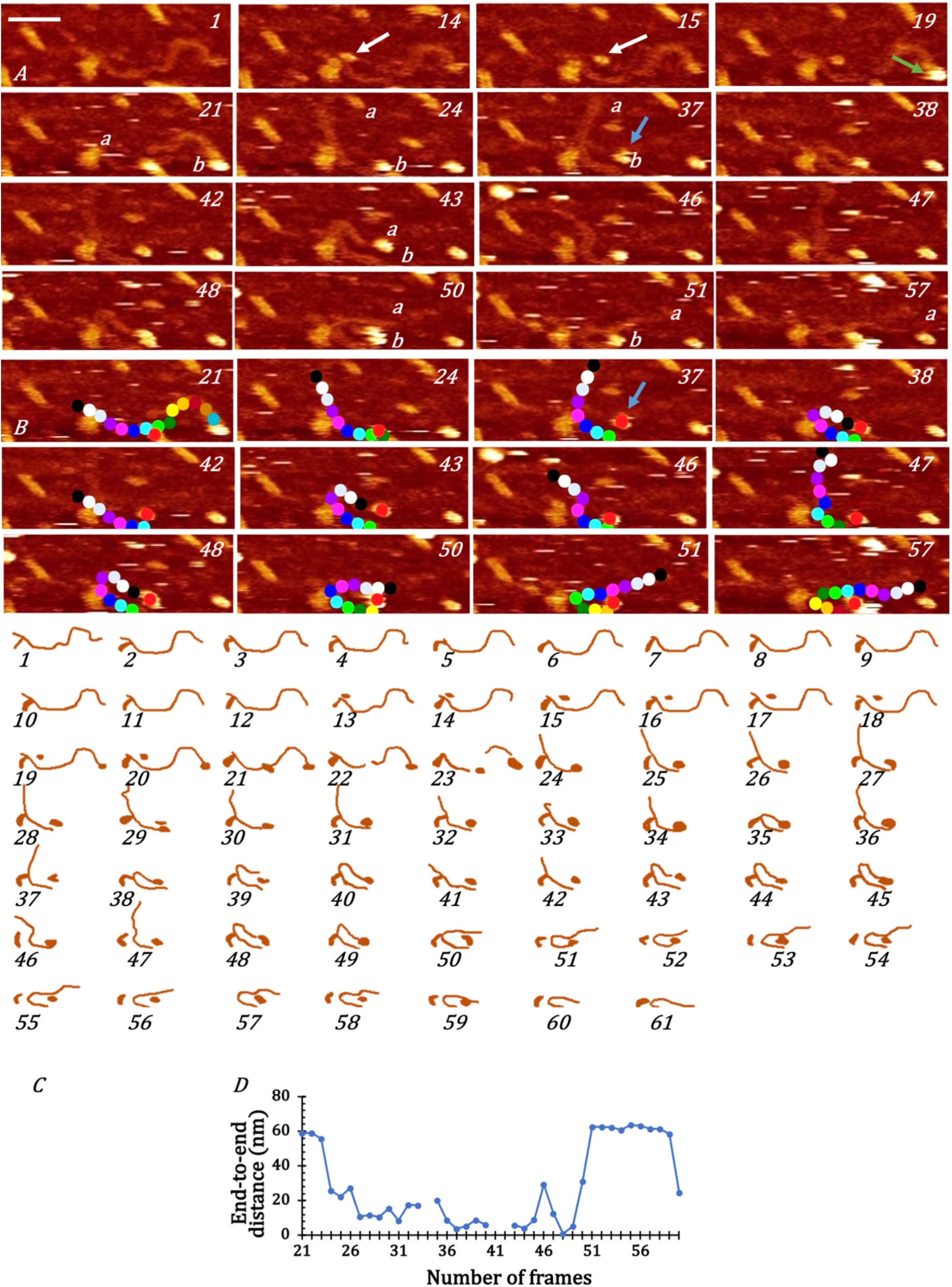
Flap DNA– T7 FEN (active) imaged in buffer (25 mM HEPES, 2 mM Mg2+, 5 mM K+) immobilized on PLO treated mica, dynamic imaging of interaction with flap DNA: height 7 nm, scale bar 50 nm, time per frame 30 s. **(A)** The frames shown here are a subset of a 79-frame scan. The protein appeared in frame 19, 17 minutes after the protein solution was added. The DNA remained in a consistent shape till 23 where the ssDNA broke and the entire strands appeared to come off and slide the dsDNA in frame 24. Refer to Movie C in Supplementary Material. **(B)** Trace of movement of segments of DNA from figure (A). The protein molecule (marked in red) could be seen bound to the end ‘b’ of the ssDNA in frame 24. **(C)** Sketch of DNA in the various frames. **(D)** End-to-end distance of DNA in the presence of active FEN protein: the DNA ends come close to one another between frames 24 and 49, which indicate the time duration the end a of ssDNA is attempting and beginning to thread, after which the ends move away as the ssDNA threading is completed.

### Protein localisation to substrate branch point is the rate limiting step during AFM imaging

From the AFM images (Figure 8A), it can be seen that the protein catalysing the breaking of the DNA strand spanned a few frames (frames 14-24 in Figure 8A) over a time frame of ∼ 5 minutes. On the contrary, the process of proteins finding and binding the DNA catalysis site took a longer time (about 15 -60 minutes, varying in the experimental repeats), partially due to the restriction of the movement of DNA as well as protein molecules due to immobilization on a 2-dimensional surface, but also because of the step of DNA substrate recognition and binding being the longer time duration step than the actual catalysis step ^94^.

### Model of DNA substrate recognition and binding by FEN

From previously reported research, the most commonly accepted theory of FEN binding is the bind then thread adaptation of the Threading model (Figure 1) according to which FEN initially recognizes the branching point ^21,40^ and not the 5’ flap end of DNA. The protein bends the DNA junction, threads 5’ flap and recognises the 3’ OH at the branch point ^20,23,25,30,31,35^. It then encloses a single 3’ nucleotide, allowing the cleaved product (resultant nicked DNA) to be ready for ligation and directs the 5’ ssDNA flap through a conserved helical arch by a threading mechanism ^15,20,24,32–35,38,43,49,95–97^. Though the results reported here support this Threading model of DNA substrate recognition by FEN, we propose the following additional steps in the recognition mechanism (Figure 9):

**Figure 9:**
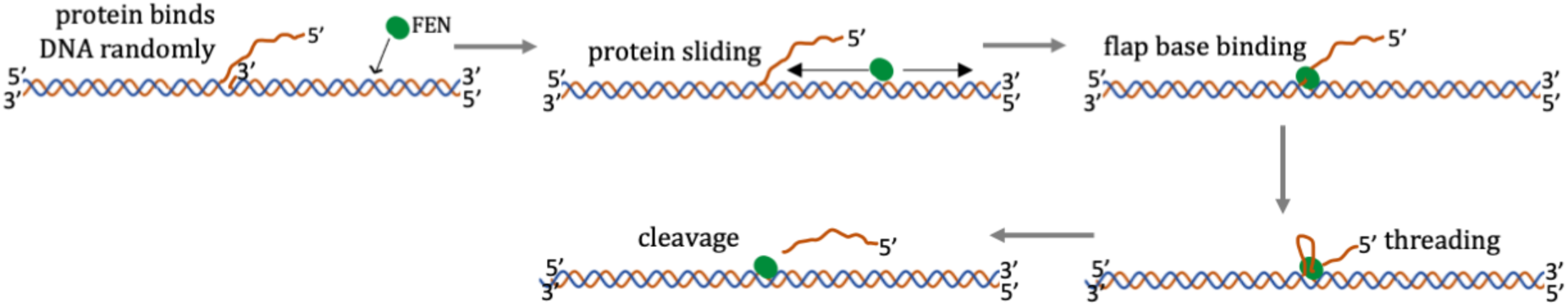
Proposed Threading model for flap DNA recognition and biding by FEN. (modification of the Threading model taken from Gloor *et al* (2010)^21^): FEN or DNAPolI binds randomly to DNA, the protein slides to find the branch point, anchors to the branch point, followed by bending and threading of the 5’ end and cleaving of the ssDNA. The diagram is not drawn to scale.

*Firstly*, the surface residues of FEN are capable of attracting and binding the negatively charged DNA anywhere along its length and threading or sliding along the length of DNA, irrespective of the position it bound to on the DNA.

*Secondly*, the process of localizing to the branch point is slower than the catalytic event of flap hydrolysis.

Thereby, in order to explain how DNA Polymerase I or FEN recognizes overhang or flap DNA for nuclease activity in the cell, it could be proposed that these proteins can randomly bind DNA at any position. This binding is followed by rapid sliding of the protein along the length of DNA, until it either falls off the substrate or encounters the bifurcation, for which it has a much higher affinity and subsequently facilitates the 5’ ssDNA bending and threading, as already proposed by the Threading model ^14^. Catalysis is limited by the encounter of the branch point, either through chance encounter of the branch point directly in the cell, or by random binding to DNA followed by sliding to reach the branch point.

## Conclusion

High-resolution AFM has been applied to image DNA Polymerase I and FEN molecules to visualize their different conformations. Mostly the protein molecules appeared globular, but on closer inspection, images of DNAPolI revealed somewhat elongated structures folded together. In some very rare cases, the protein appeared in an extended form where the domains were clearly visible: the elongated sphere-shaped FEN domain and the Klenow domain with a hand, thumb and finger shape.

In the current work, it was possible to image FEN interacting with flap or overhang DNA, causing the latter to undergo different conformational changes. A method of motion tracking and quantification of the movement of sections of DNA was devised using Adobe Illustrator, Tracker (Video Analysis and Modelling Tool) and MS Excel. It allows the segmentation of DNA and tracking of each of the segments to recognise the sections of DNA most affected by the protein.

The binding of inactive FEN domain-containing DNA Polymerase I caused the mobilisation of DNA on the mica surface, indicating that protein recognition and interaction affected the DNA conformation. The protein caused the DNA to bend in its vicinity. The time duration of the binding and the changes in DNA shape could be observed in the short time span of the protein’s appearance close to DNA. It was observed that FEN bound to DNA (both flap and overhang) randomly anywhere. The binding of the protein appeared to cause the DNA to overcome the electrostatic force of attraction binding it to the PLO treated surface, thereby indicating that the protein disrupted the charges around DNA to cause it to be attracted towards the protein. FEN was also seen sliding on the DNA and consequently bending it along its way. Based on these observations, it can be proposed that DNA Polymerase I containing FEN bind to the DNA randomly and slide on it until it encounters the branch point of ssDNA to which it anchors leading to catalysis.

Active FEN protein was observed intermittently mobilizing the DNA, sliding (and possibly threading, though this could not be observed by the limited resolution of these data) and cutting the single-stranded branch of DNA. Motion tracking of segments of DNA showed that most of the movement occurred during the short time span when the protein encountered the DNA and was restricted to the sections around the protein. It causes the single strand to bend toward the duplex region. It was interesting to observe that the protein generated an attractive interaction with the DNA that was stronger than the interaction binding the DNA to PLO treated mica and caused the DNA to mobilize and bend towards the protein. The DNA then anchored to the surface again after a few frames indicating that the effect of the protein causing DNA mobility had been removed.

As a result of the dynamic AFM imaging results obtained in the current work, the Threading model of DNA substrate recognition by FEN (that the protein bound to the branch point of DNA to perform the nuclease activity) can be corroborated with the following additional steps in the recognition mechanism: attraction and binding of the negatively charged DNA anywhere along its length by the surface residues of FEN (and DNA Polymerase I) and the sliding of the protein along the length of DNA, irrespective of the position it bound to on the DNA. Unsurprisingly, the nuclease activity is the faster step in the overall catalytic cycle, as the enzyme activity is limited by the rate of encountering the branch point, either randomly in the cell, or by random binding to DNA followed by sliding to reach the branch point.

Future experiments could be performed with improved spatial and time resolution AFM to observe the conformation changes and the mechanism of catalysis, *e.g.* the modification of the structure of FEN active site during the reaction and the changes on the surface residues of FEN that allow it to bend the ssDNA to thread through. The importance of various mutations in the protein and their consequences in disorders like cancer could be studied by dynamic AFM imaging of the mutants interacting with DNA. These experiments could be expanded to understand not only the functioning of FEN, but the entire DNA Polymerase I in the highly regulated process of DNA replication and cell division.

## Supporting information

Supplemental Movie A for Figure 6

Supplemental Movie B for Figure 7

Supplemental Movie C for Figure 8

## DATA AVAILABILITY

The data underpinning this article are available in the article and in the online supplementary material.

## SUPPLEMENTARY DATA

Movie A for Figure 6, Movie B for Figure 7, Movie C for Figure 8 and Supplementary Information containing supplementary Tables and Figures from the Supplementary Material link.

## ACKNOWLEDGEMENTS

We are indebted to Drs Laia Pasquina Lemonche and Nic Mullin for help with AFM.

## Author contributions

V.V. performed all AFM experiments and analyses with input from J.K.H. The manuscript and figures were drafted by V.V., E.L.B., J.R.S and J.K.H. Protein expression, characterisation and cloning work was carried out by E.L.B., S.C., S.C.K. and V.V. with J.P. providing technical support. J.K.H. and J.R.S. conceived the project and experimental design, provided supervision, and obtained funding. All authors approved the final version of the manuscript.

