## Supplementary Tables and Figures for "Flap endonuclease Substrate Interactions studied using Dynamic Single-Molecule Atomic Force Microscopy"

**Table of Contents**

**Supplementary Methods** **Page 2**

**Supplementary Figures S1 – S3** **Pages 3–5**

**Supplementary Table S1** **Page 6**

**Supplementary Information – gene fragments** **Page 7–8**

**Supplementary Videos** **Page 9**

### Supplementary Methods

#### Zymogram for T7FEN

A 10% SDS-PAGE gel was cast with 300 μg type XIV DNA (Sigma, in 25 mM Tris-HCl, pH 8) and samples were prepared and run as for a regular SDS-PAGE gel. After electrophoresis, the gel was washed (3 X 20 minutes) with Tris-Bicine-glycerol (TBG) buffer (100 mM Tris, 100 mM Bicine, 10% (v/v) glycerol) and incubated in a zymogram reaction buffer (100 mM Tris, 100 mM Bicine, pH 8, 10% (v/v) glycerol, 100 mM KCl, 50 mM NaCl, 10 mM $MgCl_2$, and 1 mM DTT) at room temperature for 15 minutes. The gel was stained with 0.5 $\mu L.mL^{-1}$ Midori Green Advance stain and imaged using the BIO-RAD EZ gel documentation system under the SYBR green setting. Nuclease activity was visualised as the presence of a dark band.

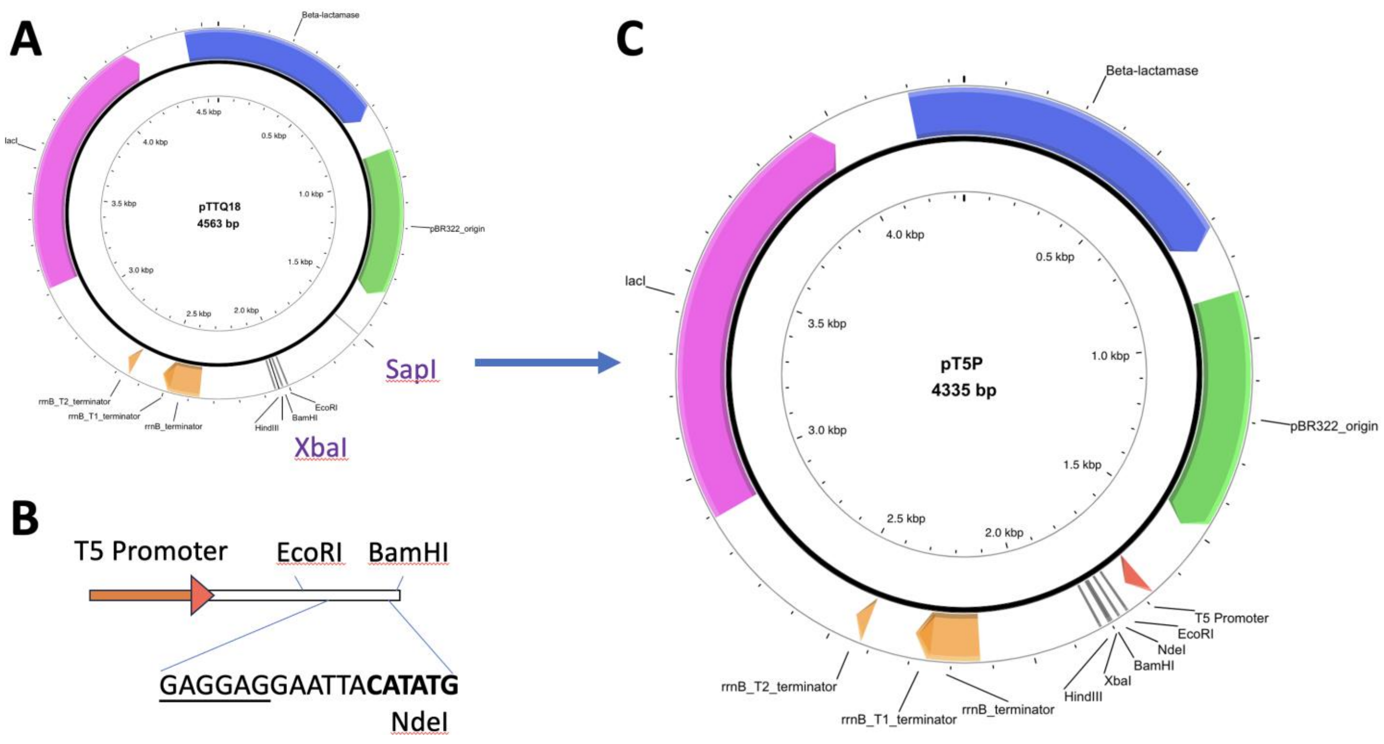


**Figure S1 Schematic diagram showing construction and features of expression plasmid pT5P**

Plasmid pTTQ18 **(A)** was digested with SapI and XbaI, and ligated with synthetic oligonucleotides **(B)** which encode a bacteriophage T5 promoter controlled by lacI operator sequences. It also includes a ribosomal binding site (underlined), start codon and in-frame NdeI site (bold). **(C)** Map of resulting plasmid pT5P, the SapI site is lost during this process.

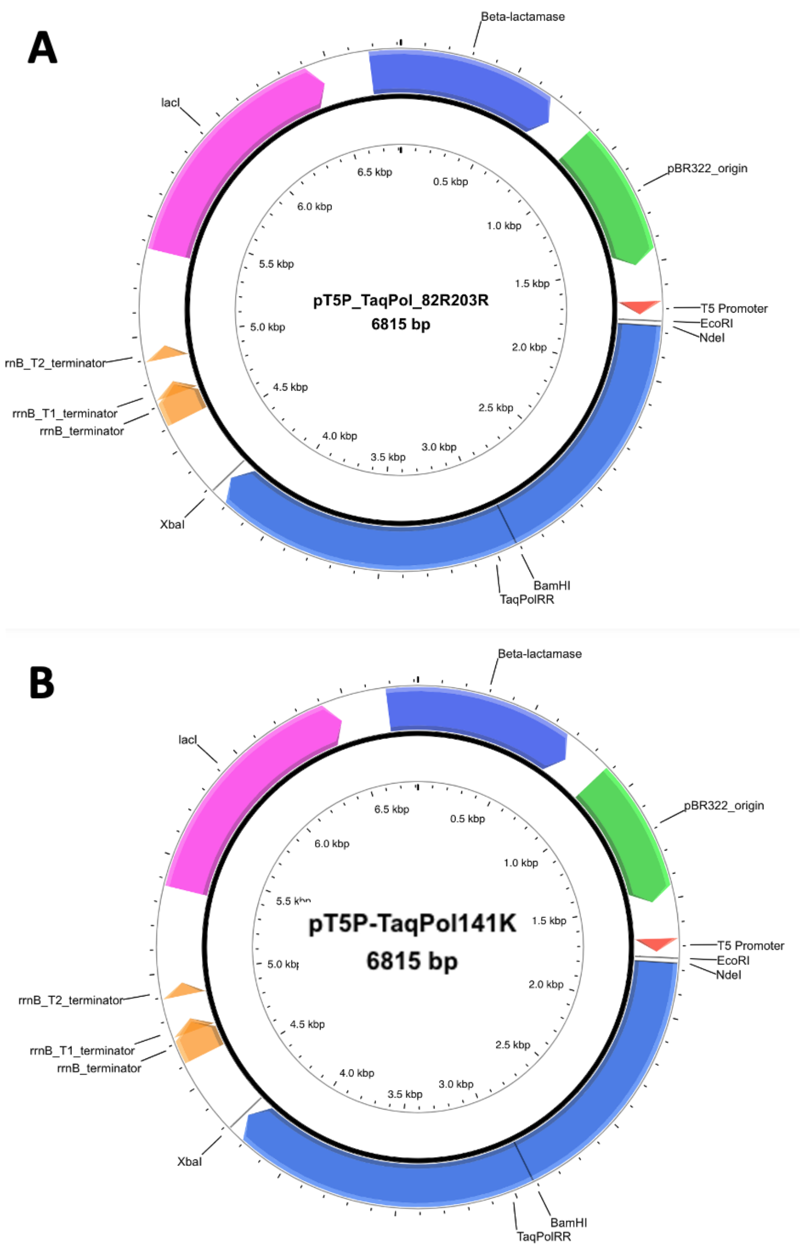


**Figure S2 Construction of Taq Polymerase expression plasmids**

**(A)** Schematic diagram showing the plasmid pT5P-TaqPol82-203RR. **(B)** The synthetic EcoRI-BamHI fragment reverting the Arg82 and Arg203 back to Lys82 and Thr203 and introducing the Asp141Lys mutation was used to replace the equivalent fragment in pT5P-TaqPol82-203RR to yield pT5P-TaqPol141K.

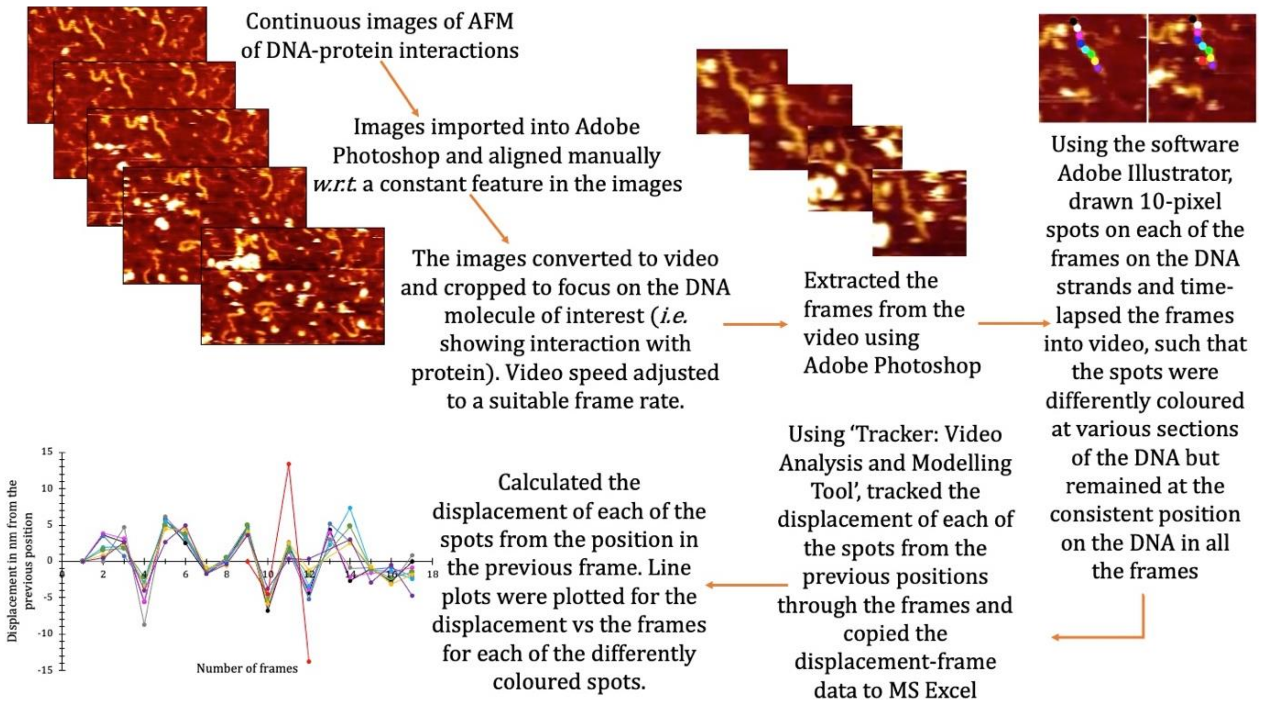


**Figure S3 Methodology for DNA Motion Tracking**
Outline of the image processing into videos for tracking the movement of sections of DNA under the influence of the protein.

### Table S1 Oligonucleotides used in this study

| Name (Length) | Sequence (5′–3′) |
|---|---|
| **Oligo 1** (79) | *CGGGTGTTGGCGGGTGTCGGGGCTGGCTTCTGCTCTTCCCACGAATCGATCACGCCACTTCTCCA TTGGTGCCAAATTA |
| **Oligo 2** (150) | $(T)_{128}$GCCTTTTGTTTCTTGGTGTATG |
| **Oligo 3** (120) | *GAGACTCGCTTCTTTTGAAATGCATTGATCGCGCGTTTCGGTGATGACGGTGAAAACCTCTGACA CATGCAGCTCCCGGACCTGCGGTTACCCATGGCCTGTAATCCAGCTCGAGTCAAG |
| **Oligo 4** (80) | *AAGTGGCGTGATCGATTCGTGGGAAGAGCAGAAGCCAGCCCCGACACCCGCCAACACCCGCTGAC GCGCCCTGACGGGCT |
| **Oligo 5** (33) | *AAGAAACAAAAGGCTAATTTGGCACCAATGGAG |
| **Oligo 6** (128) | CTTGACTCGAGCTGGATTACAGGCCATGGGTAACCGCAGGTCCGGGAGCTGCATGTGTCAGAGGT TTTCACCGTCATCACCGAAACGCGCGATCAATGCATTTCAAAAGAAGCGAGTCTCCATACACC |
| **Oligo 8** (99) | ATTGTACTGAGAGTGCACCATATGCGGTGTGAAATACCGCACAGATGCGTAAGGAGAAAGCGGAT GCCGGGAGCAGACAAGCCCGTCAGGGCGCGTCAG |
| **Oligo 9** (79) | *TGTCTGCTCCCGGCATCCGCTTTCTCCTTACGCATCTGTGCGGTATTTCACACCGCATATGGTGC ACTCTCAGTACAAT |
| **T5Pr01** (48) | AGCCTCGAGAAATCATAAAAAATTTATTTGCTTTGTGAGCGGATAACA |
| **T5Pr02** (49) | ATTATAATAGATTCAATTG TGAGCGGATAACAATTTC ACACAGAATTCA |
| **T5Pr03** (45) | TTAAA<u>GAGGAG</u>GAATTA**CATATG**GAGCTCGGTACCCGGGGATCCT |
| **T5Pr04** (24) | CTAGAGGATCCCCGGGTACCGAGC |
| **T5Pr05** (38) | TCCATATGTAATTCCTCCTCTTTAATGAATTCTGTGTG |
| **T5Pr06** (45) | AAATTGTTATCCGCTCAC AATTGAATCTATTATAATT GTTATCCG |
| **T5Pr07** (37) | CTCACAAAGCAAATAAAT TTTTTATGATTTCTCGAG |
| **T7exoF** (22) | AGCTGACGAATACGTAGCGGAA |
| **T7exoR** (34) | AAAACTGCAGTTACTACGGTCTCCACAGGTAAAT |
| * Indicates 5′-phosphorylated oligonucleotide. Ribosomal binding site underlined and NdeI site CATATG in bold in T5Pr03. | |

### Sequences of gene fragments used in this study

**>Construct 1:** Synthetic fragment encoding Thermus aquaticus DNA polymerase I mutant (Lys82Arg; Thr203Arg). Start and stop codons highlighted in yellow and the unique internal BamHI site used for producing the wild-type coding sequence and TaqFEN domain is shown highlighted in cyan.

GAATTCTAAAAAGGAGGAAAACAT**ATG**GGTATGCTGCCGCTGTTCGAGCCGAAGGGTCGTGTGCTGCTGGTTGACGGCCACCACCTGGCG
TACCGTACCTTTCACGCGCTGAAGGGTCTGACCACCAGCCGTGGTGAACCGGTGCAGGCGGTTTATGGTTTCGCGAAAAGCCTGCTGAAG
GCGCTGAAAGAGGACGGCGATGCGGTGATCGTGGTTTTCGATGCGAAGGCGCCGAGCTTTCGTCACGAAGCGTACGGTGGCTATCGTGCG
GGTCGTGCGCCGACCCCGGAGGACTTTCCGCGTCAACTGGCGCTGATTAAAGAACTGGTTGATCTGCTGGGTCTGGCGCGTCTGGAAGTG
CCGGGCTACGAAGCGGACGATGTTCTGGCGAGCCTGGCGAAGAAAGCGGAGAAGGAAGGTTACGAAGTGCGTATCCTGACCGCGGACAAG
GACCTGTATCAGCTGCTGAGCGACCGTATCCACGTTCTGCACCCGGAGGGTTATCTGATTACCCCGGCGTGGCTGTGGGAAAAGTACGGC
CTGCGTCCGGACCAATGGGCGGATTATCGTGCGCTGACCGGTGACGAGAGCGATAACCTGCCGGGCGTGAAAGGTATTGGCGAAAAACGT
GCGCGTAAGCTGCTGGAGGAATGGGGTAGCCTGGAAGCGCTGCTGAAAAACCTGGATCGTCTGAAGCCGGCGATCCGTGAGAAAATTCTG
GCGCACATGGACGATCTGAAGCTGAGCTGGGACCTGGCGAAAGTGCGTACCGACCTGCCGCTGGAAGTGGACTTCGCGAAGCGTCGTGAG
CCGGATCGTGAACGTCTGCGTGCGTTCCTGGAGCGTCTGGAATTTGGTAGCCTGCTGCACGAGTTTGGCCTGCTGGAAAGCCCGAAGGCG
CTGGAGGAAGCGCCGTGGCCGCCGCCAGAGGGTGCGTTCGTGGGCTTTGTTCTGAGCCGTAAAGAACCGATGTGGGCGGACCTGCTGGCG
CTGGCGGCGGCGCGTGGTGGCCGTGTGCACCGTGCGCCGGAGCCGTACAAGGCGCTGCGTGACCTGAAAGAAGCGCGTGGTCTGCTGGCG
AAGGACCTGAGCGTTCTGGCGCTGCGCGAAGGTCTGGGTCTGCCGCCGGGTGACGATCCGATGCTGCTGGCGTACCTGCT**GGATCC**GAGC
AACACCACCCCGGAGGGTGTGGCGCGTCGTTATGGTGGCGAATGGACCGAGGAAGCGGGCGAGCGTGCGGCGCTGAGCGAACGTCTGTTC
GCGAACCTGTGGGGTCGTCTGGAGGGCGAGGAACGTCTGCTGTGGCTGTACCGTGAGGTGGAACGTCCGCTGAGCGCGGTTCTGGCGCAC
ATGGAAGCGACCGGTGTGCGTCTGGACGTTGCGTATCTGCGTGCGCTGAGCCTGGAAGTGGCGGAGGAAATCGCGCGTCTGGAGGCGGAA
GTTTTCCGTCTGGCGGGCCACCCGTTTAACCTGAACAGCCGTGACCAGCTGGAGCGTGTTCTGTTTGATGAACTGGGTCTGCCGGCGATTG
GCAAGACCGAAAAAACCGGCAAGCGTAGCACCAGCGCGGCGGTGCTGGAGGCGCTGCGTGAAGCGCACCCGATCGTTGAGAAGATTCTGC
AGTACCGTGAACTGACCAAGCTGAAAAGCACCTATATCGACCCGCTGCCGGATCTGATTCACCCGCGTACCGGTCGTCTGCACACCCGTTT
CAACCAAACCGCGACCGCGACCGGCCGTCTGAGCAGCAGCGACCCGAACCTGCAGAACATCCCGGTGCGTACCCCGCTGGGTCAACGTATC
CGTCGTGCGTTTATTGCGGAGGAAGGCTGGCTGCTGGTTGCGCTGGATTACAGCCAGATTGAGCTGCGTGTTCTGGCGCACCTGAGCGGT
GACGAAAACCTGATCCGTGTTTTCCAAGAGGGCCGTGATATTCACACCGAAACCGCGAGCTGGATGTTTGGTGTGCCGCGTGAGGCGGTT
GACCCGCTGATGCGTCGTGCGGCGAAGACCATCAACTTCGGTGTGCTGTATGGCATGAGCGCGCACCGTCTGAGCCAGGAGCTGGCGATC
CCGTACGAGGAAGCGCAGGCGTTTATTGAACGTTATTTCCAAAGCTTTCCGAAGGTTCGTGCGTGGATTGAGAAAACCCTGGAGGAAGGT
CGTCGTCGTGGTTACGTGGAAACCCTGTTCGGTCGTCGTCGTTACGTTCCGGATCTGGAGGCGCGTGTGAAAAGCGTTCGTGAGGCGGCG
GAACGTATGGCGTTCAACATGCCGGTGCAGGGTACCGCGGCGGACCTGATGAAACTGGCGATGGTTAAGCTGTTTCCGCGTCTGGAGGAA
ATGGGCGCGCGTATGCTGCTGCAAGTGCACGATGAGCTGGTTCTGGAAGCGCCGAAGGAGCGTGCGGAAGCGGTGGCGCGTCTGGCGAAA
GAGGTGATGGAAGGTGTTTACCCGCTGGCGGTTCCGCTGGAAGTTGAGGTGGGTATCGGCGAGGACTGGCTGAGCGCGAAAGAA**TAA**TA
ATAAGATCT

**>Construct 2:** Synthetic fragment encoding *Thermus aquaticus* DNA polymerase I mutant (Asp141Lys, codon highlighted in green). Start and stop codons highlighted in yellow and the unique internal BamHI site used for cloning is highlighted in cyan.

GAATTCTAAAAAGGAGGAAAACAT**ATG**GGTATGCTGCCGCTGTTCGAGCCGAAGGGTCGTGTGCTGCTGGTTGACGGCCACCACCTGGCG
TACCGTACCTTTCACGCGCTGAAGGGTCTGACCACCAGCCGTGGTGAACCGGTGCAGGCGGTTTATGGTTTCGCGAAAAGCCTGCTGAAG
GCGCTGAAAGAGGACGGCGATGCGGTGATCGTGGTTTTCGATGCGAAGGCGCCGAGCTTTCGTCACGAAGCGTACGGTGGCTATAAAGCG
GGTCGTGCGCCGACCCCGGAGGACTTTCCGCGTCAACTGGCGCTGATTAAAGAACTGGTTGATCTGCTGGGTCTGGCGCGTCTGGAAGTG
CCGGGCTACGAAGCGGACGATGTTCTGGCGAGCCTGGCGAAGAAAGCGGAGAAGGAAGGTTACGAAGTGCGTATCCTGACCGCGAAAAA
GGACCTGTATCAGCTGCTGAGCGACCGTATCCACGTTCTGCACCCGGAGGGTTATCTGATTACCCCGGCGTGGCTGTGGGAAAAGTACGG
CCTGCGTCCGGACCAATGGGCGGATTATCGTGCGCTGACCGGTGACGAGAGCGATAACCTGCCGGGCGTGAAAGGTATTGGCGAAAAAAC
TGCGCGTAAGCTGCTGGAGGAATGGGGTAGCCTGGAAGCGCTGCTGAAAAACCTGGATCGTCTGAAGCCGGCGATCCGTGAGAAAATTCT
GGCGCACATGGACGATCTGAAGCTGAGCTGGGACCTGGCGAAAGTGCGTACCGACCTGCCGCTGGAAGTGGACTTCGCGAAGCGTCGTGA
GCCGGATCGTGAACGTCTGCGTGCGTTCCTGGAGCGTCTGGAATTTGGTAGCCTGCTGCACGAGTTTGGCCTGCTGGAAAGCCCGAAGGC
GCTGGAGGAAGCGCCGTGGCCGCCGCCAGAGGGTGCGTTCGTGGGCTTTGTTCTGAGCCGTAAAGAACCGATGTGGGCGGACCTGCTGGC
GCTGGCGGCGGCGCGTGGTGGCCGTGTGCACCGTGCGCCGGAGCCGTACAAGGCGCTGCGTGACCTGAAAGAAGCGCGTGGTCTGCTGGC
GAAGGACCTGAGCGTTCTGGCGCTGCGCGAAGGTCTGGGTCTGCCGCCGGGTGACGATCCGATGCTGCTGGCGTACCTGCT**GGATCC**GAG
CAACACCACCCCGGAGGGTGTGGCGCGTCGTTATGGTGGCGAATGGACCGAGGAAGCGGGCGAGCGTGCGGCGCTGAGCGAACGTCTGTT
CGCGAACCTGTGGGGTCGTCTGGAGGGCGAGGAACGTCTGCTGTGGCTGTACCGTGAGGTGGAACGTCCGCTGAGCGCGGTTCTGGCGCA
CATGGAAGCGACCGGTGTGCGTCTGGACGTTGCGTATCTGCGTGCGCTGAGCCTGGAAGTGGCGGAGGAAATCGCGCGTCTGGAGGCGGA
AGTTTTCCGTCTGGCGGGCCACCCGTTTAACCTGAACAGCCGTGACCAGCTGGAGCGTGTTCTGTTTGATGAACTGGGTCTGCCGGCGAT
TGGCAAGACCGAAAAAACCGGCAAGCGTAGCACCAGCGCGGCGGTGCTGGAGGCGCTGCGTGAAGCGCACCCGATCGTTGAGAAGATTCT
GCAGTACCGTGAACTGACCAAGCTGAAAAGCACCTATATCGACCCGCTGCCGGATCTGATTCACCCGCGTACCGGTCGTCTGCACACCCGT
TTCAACCAAACCGCGACCGCGACCGGCCGTCTGAGCAGCAGCGACCCGAACCTGCAGAACATCCCGGTGCGTACCCCGCTGGGTCAACGTA
TCCGTCGTGCGTTTATTGCGGAGGAAGGCTGGCTGCTGGTTGCGCTGGATTACAGCCAGATTGAGCTGCGTGTTCTGGCGCACCTGAGCG
GTGACGAAAACCTGATCCGTGTTTTCCAAGAGGGCCGTGATATTCACACCGAAACCGCGAGCTGGATGTTTGGTGTGCCGCGTGAGGCGG
TTGACCCGCTGATGCGTCGTGCGGCGAAGACCATCAACTTCGGTGTGCTGTATGGCATGAGCGCGCACCGTCTGAGCCAGGAGCTGGCGA
TCCCGTACGAGGAAGCGCAGGCGTTTATTGAACGTTATTTCCAAAGCTTTCCGAAGGTTCGTGCGTGGATTGAGAAAACCCTGGAGGAAG
GTCGTCGTCGTGGTTACGTGGAAACCCTGTTCGGTCGTCGTCGTTACGTTCCGGATCTGGAGGCGCGTGTGAAAAGCGTTCGTGAGGCGG
CGGAACGTATGGCGTTCAACATGCCGGTGCAGGGTACCGCGGCGGACCTGATGAAACTGGCGATGGTTAAGCTGTTTCCGCGTCTGGAGG
AAATGGGCGCGCGTATGCTGCTGCAAGTGCACGATGAGCTGGTTCTGGAAGCGCCGAAGGAGCGTGCGGAAGCGGTGGCGCGTCTGGCGA
AAGAGGTGATGGAAGGTGTTTACCCGCTGGCGGTTCCGCTGGAAGTTGAGGTGGGTATCGGCGAGGACTGGCTGAGCGCGAAAGAA**TAA**
TAATAAGATCT

1. Movie A for Figure 6
2. Movie B for Figure 7
3. Movie C for Figure 8
